## Supplementary Material for "Whole-genomes from the extinct Xerces Blue butterfly can help identify declining insect species"

### Supplementary Information

This file includes:

Materials and Methods

Figs. S1 to S11

Tables S1 to S6

References

### Supplementary Materials and Methods

#### Software information

| Software | Version | Source |
|---|---|---|
| *BUSCO* | *v.5.1.2* | 1 |
| *AdapterRemoval* | *v.2.2.2* | 2 |
| *BWA – backtrack* | *v.0.7.1* | 3 |
| *BWA – mem* | *v.0.7.1* | 4 |
| *Qualimap2* | *v.2.2.2* | 5 |
| *pmdtools* | *v.0.50* | 6 |
| *MapDamage2* | *v.2.7.12* | 7 |
| *Bedtools* | *v.2.27.1* | 8 |
| *snpAD* | *v.0.3.2* | 9 |
| *GATK* | *v.3.5 – 3.7* | 10 |
| *vcftools* | *v.0.1.12b – 0.1.14b* | 11 |
| *angsd* | *v.0.916* | 12 |
| *bcftools* | *v.1.9* | 13 |
| *Mitofinder* | v.1.4 | 14 |
| *MACSE* | v.2.05 | 15 |

| | | |
|---|---|---|
| *MAFFT* | v.7.490 | 16 |
| *IQ-TREE2* | v.2.1.3 | 17 |
| *ModelFinder* | *Avail. In IQ-TREE2* | 18 |
| *UFBoot2* | *Avail. In IQ-TREE2* | 19 |
| *BEAST2* | v.2.6.3 | 20 |
| bModelTest | v.1.2.1 | 21 |
| *Tracer* | v.1.7.2 | 22 |
| *PSMC* | *v.0.6.5* | 23 |
| PCAngsd | v.20180209 | 24 |
| *Bcftools-roh* | *v.1.9* | 25 |
| *SNPeff* | *v.4.3* | 26 |
| *Picard* | v.2.0.1 | 27 |
| *Samtools* | v.1.6 | 28 |
| *BamUtil* | v.1.0.13 | 29 |
| *Bedtools* | v.2.27.1 | 8 |
| *BLAST* | v.2.2.2 | 30 |
| *BBMap* | *v.38.18* | 31 |
| *Prinseq* | v.0.20.4 | 32 |
| *Kraken2* | v.2.1.1 | 33 |
| *R* | *v.3.6.3 – 4.1.0* | 34 |
| *Ggplot2* | v.3.0.0 | 35 |

**DNA extraction and sequencing**

One ml of digestion buffer (final concentrations: 3 mM CaCl2, % SDS, 40 mM DTT, 0.25 mg/ml proteinase K, 100 mM Tris buffer pH 8.0 and 100 mM NaCl) was added to each crushed butterfly residue, including an extraction blank, and incubated at 37 °C overnight (24h) on rotation (750-900 rpm). Next, DNA extraction was continued following the method proposed by Dabney et al. 2013. Remaining butterfly sample was then pelleted by centrifugation in a bench-top centrifuge for 2 min at maximum speed (16,100 × g). The supernatant was added to 10 mL of binding buffer (final concentrations: 5 M guanidine hydrochloride, 40% (vol/vol) isopropanol, 0.05% Tween-20, and 90 mM sodium acetate (pH 5.2)) and purified on a High Pure Extender column (Roche). DNA extracts were eluted with 45 µL of low EDTA TE buffer (pH 8.0) and quantified using a Qubit instrument.

Following extraction, the DNA extract was converted into Illumina sequencing libraries following the BEST protocol [37]. Each library was amplified by PCR using two uniquely barcoded primers, prior to being purified with a 1.5x AMPure clean (Beckman Coulter) and eluted in 25 μl of low EDTA TE buffer (pH 8.0). One Xerces Blue sample did not yield detectable DNA in two independent extractions. For each of the successful extracts we prepared a single library which was shotgun sequenced on the HiseqX Illumina platform.

**aDNA sequences mapping**

In order to minimise the effect of aDNA post-mortem damage, which could hamper the recovery of authentic but severely damaged sequences, we use a set of mapping parameters optimised for aDNA data [38]. Instead of the commonly used BWA v.0.7.1 mem algorithm (local alignment, standard for modern DNA), we used the *backtrack* algorithm or global algorithm [3]. In short, by completely disabling the seeding length (using an arbitrarily long read length, -l 1024), the algorithm has to map the whole read against the reference genome. In addition to that, and to account for the deamination at reads' ends, we reduce the mapping astringency by reducing the edit distance value so more genetically distant sequences, or in this case damaged, will be mapped (-n 0.01). Finally, the gap open penalty value was set to 2. After mapping, duplicated reads were removed using picard MarkDuplicates. Mapped reads with mapping quality below 30 were removed using samtools. Finally, to avoid problems in the next steps derived from spurious callings due to aDNA at reads' ends, we trimmed 2 nt from each read end using BamUtil trimbam.

We mapped 124,101,622 and 184,084,237 unique DNA reads of Xerces Blue and Silvery Blue, respectively, against the *G. alexis* reference genome (Table S2). The DNA reads exhibited typical ancient DNA features, such as short mean read length (ranging from 47.55 to 67.41 bases on average, depending on the specimen (Fig. S2)) and post-mortem deamination patterns at the 5' and 3' ends (Table S2). We also have displayed the depth histogram of the samples (Fig. S09), the mapped read length distribution (Fig. S10), and the edit distance distribution (Fig. S11). The historical genomes covered 49.3% (Xerces Blue) and 55.2% (Silvery Blue) of the *G. alexis* reference genome. To estimate the mappable fraction of this reference, we randomly fragmented it to 50 to 70 nucleotides and mapped the generated fragments back to the complete genome. An average of 57.8% of the *G. alexis* genome was covered with these read lengths (Table S2). We suggest that reduced coverage from the historical specimens may be due to genomic divergence of *G. xerces* and *G. lygdamus* from the *G. alexis* reference and the presence of unmappable, repetitive regions (Fig. S2-S4) (Table S2).

**Mapping of RVcoll10-B005**

Since RVcoll10-B005 is a modern individual, we proceed to map it against the *G. alexis* reference genome with slightly altered parameters. As with the historical samples, pair were collapsed using AdapterRemoval2. BWA mem with default parameters was used as the mapping algorithm. As with the other samples, reads were filtered with Samtools (min. quality of 30) and duplicates were removed by coordinate with picard.

**Coloration genes variability**

To find possible amino acid-changing variants that could explain phenotypical differences between *G. lygdamus* and *G. xerces*, we have identified and explored three well-known genes associated to colour patterns in butterflies: *optix*, *cortex* and *Wnt* genes [39–43]. First, we located those genes in our annotation with BLAST and their homologs in other butterfly species, setting an E-value lower than 0.001 and an Identity value above 60% (Table S5). Then, the coordinates were called using GATK UnifiedGenotyper. Variants were filtered for indels and minimum Genotype Quality of 30 using. Variants were kept regardless of their coverage. A variant is considered as fixed in one species if it is covered in at least 2 individuals of each species, it is in homozygous state, and when one of the species present all their genotypes calls as homozygous for the alternative allele while in the other are homozygous of the reference allele. No fixated mutations were identified in the regions covered at the same time by *G. lygdamus* and *G. xerces* sequences.

### Supplementary Figures

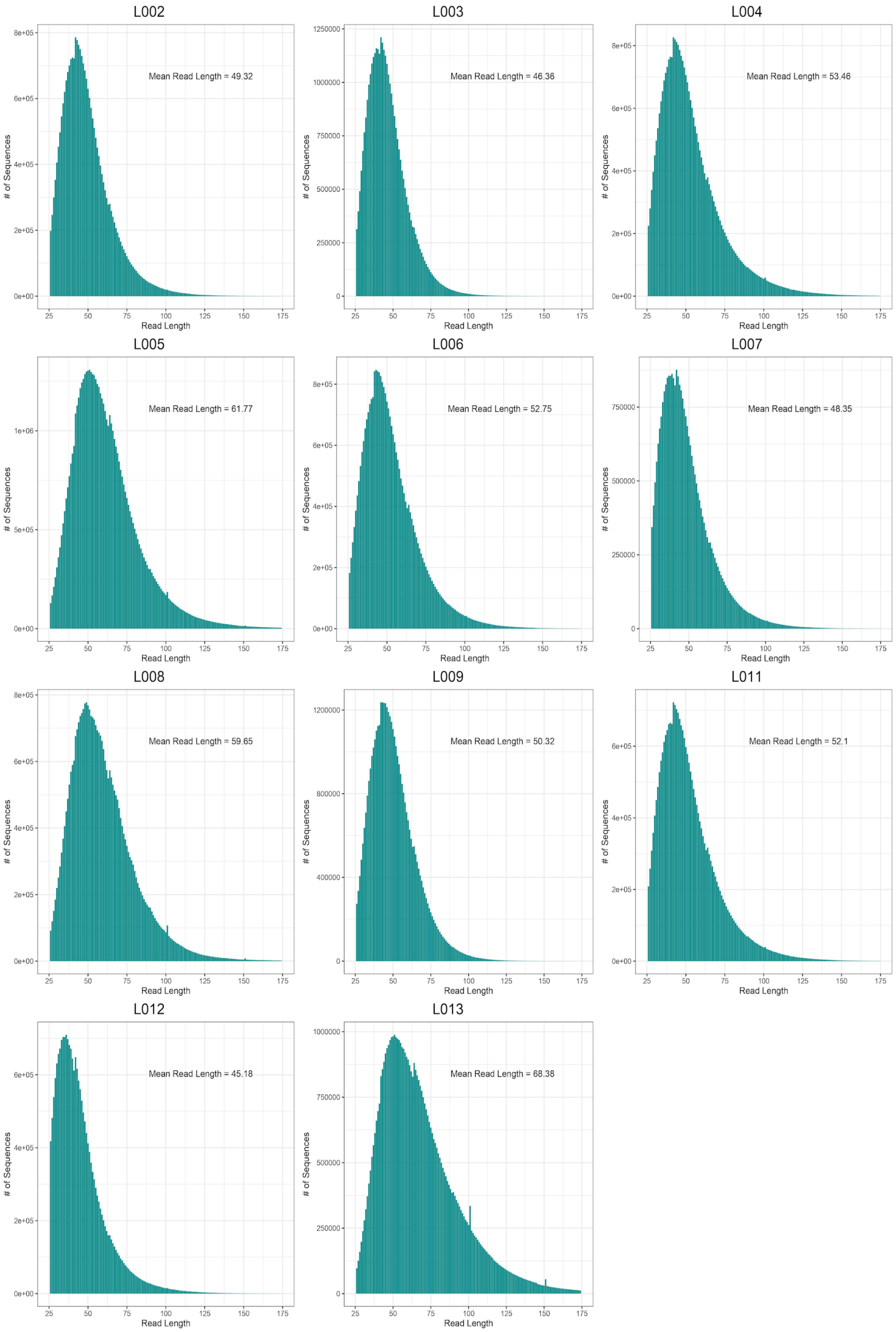


**Fig. S1: Read length distribution across the 11 analysed historical samples.**

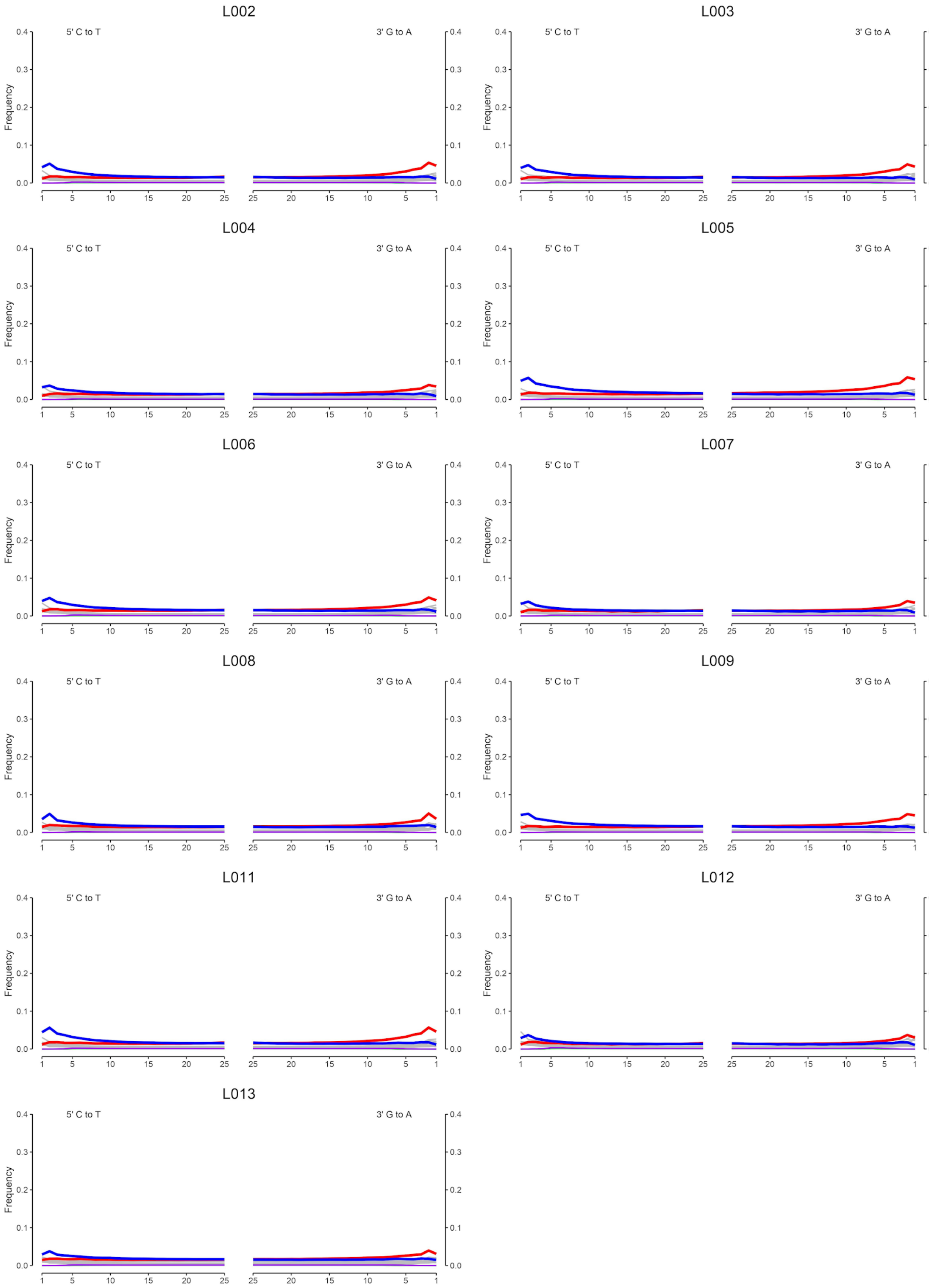


**Fig. S2: Post-mortem damage patterns of the analysed historical samples.** Frequency of C to T (blue) and G to A (red) substitutions are displayed across the last 25 bases of each end.

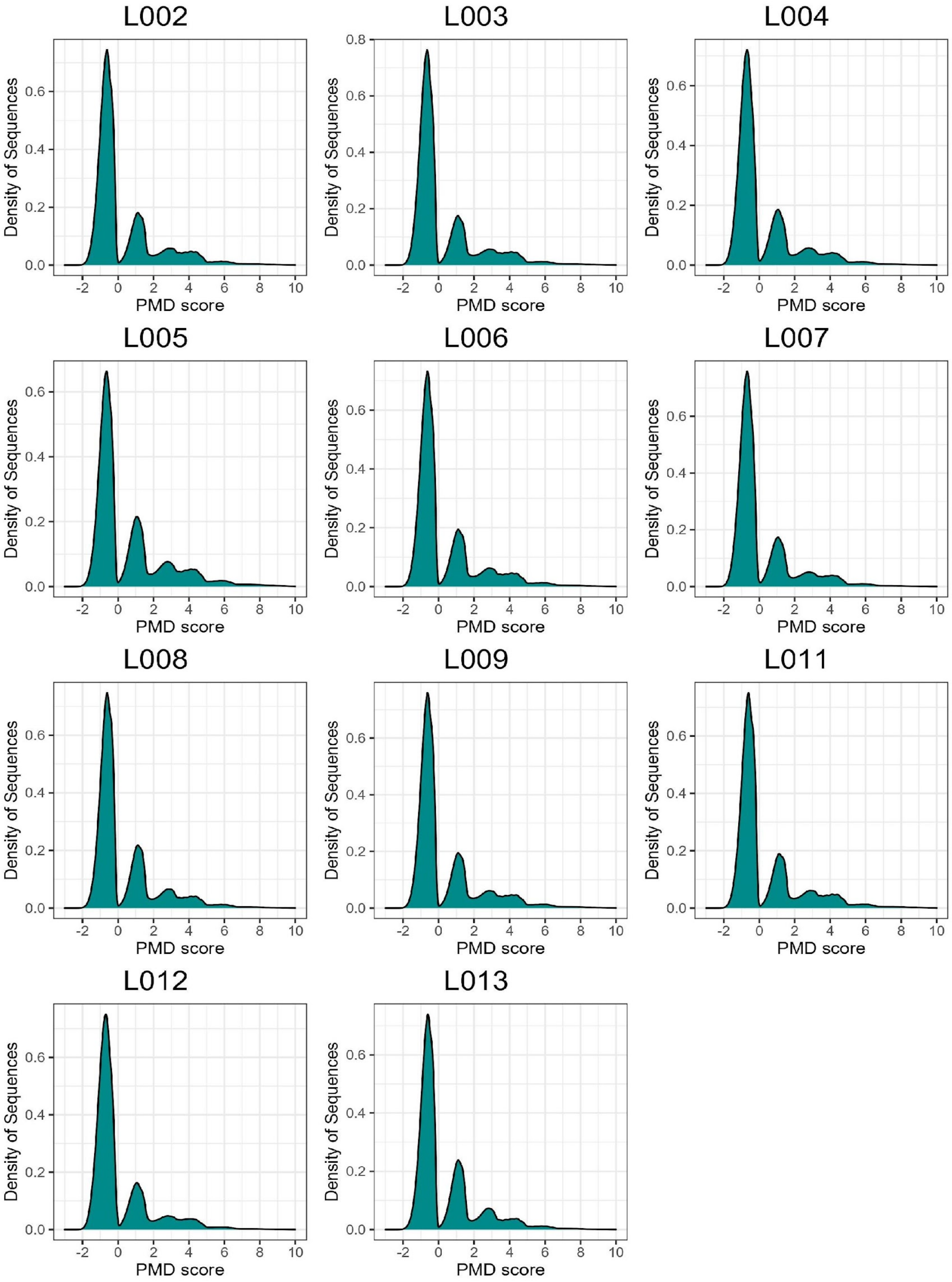


**Fig. S3: PMD score distribution of the analysed historical samples.**

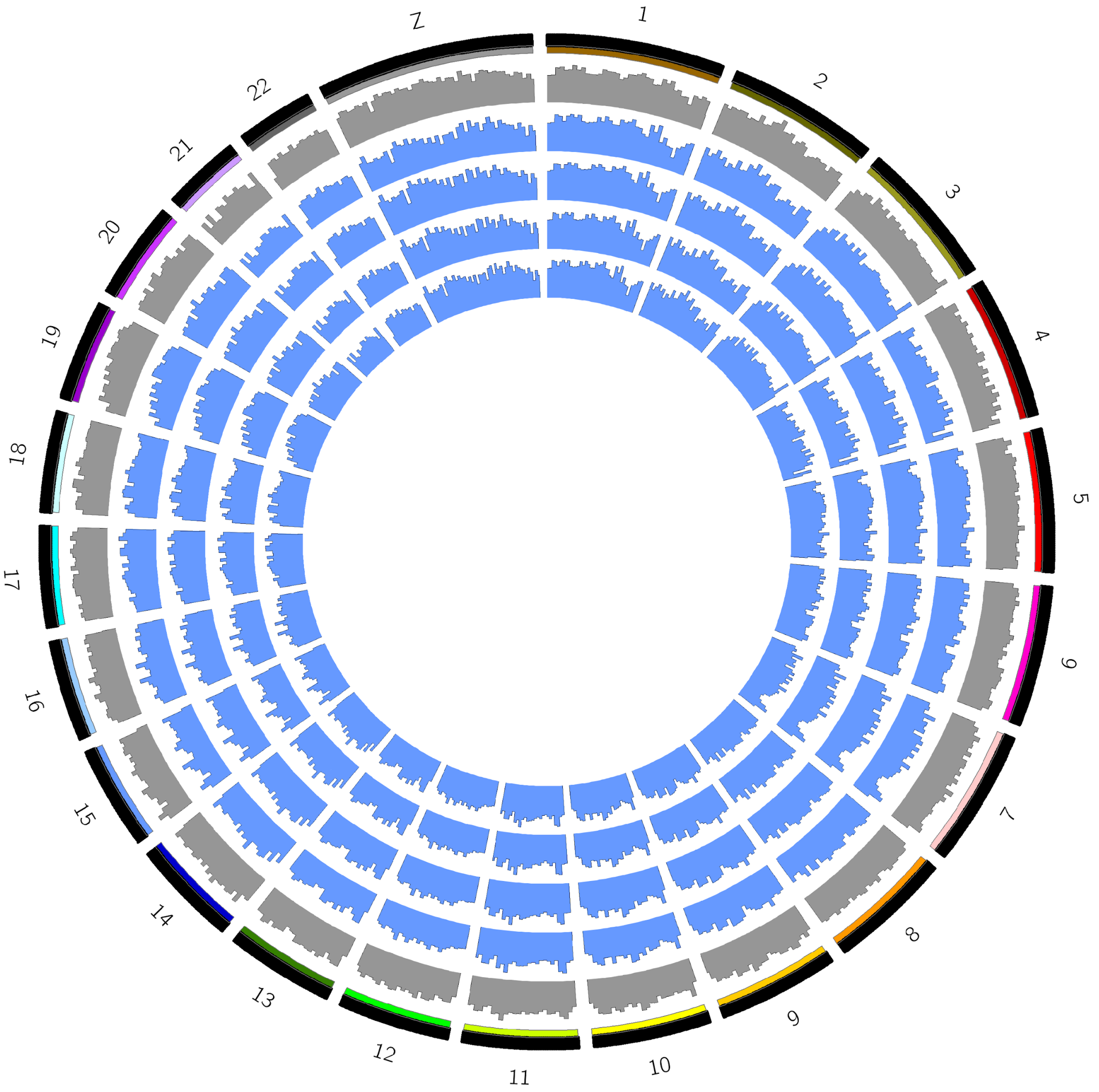


**Fig. S4: Coverage distribution of *G. xerces* samples across the reference.** Reference mappability is represented by the outermost grey histogram. Blue histograms represent the coverage of *G. xerces* samples across the *G. alexis* reference in windows of 1mbp. The sample represented are, from outer to inner rings: L003, L005, L007 and L009.

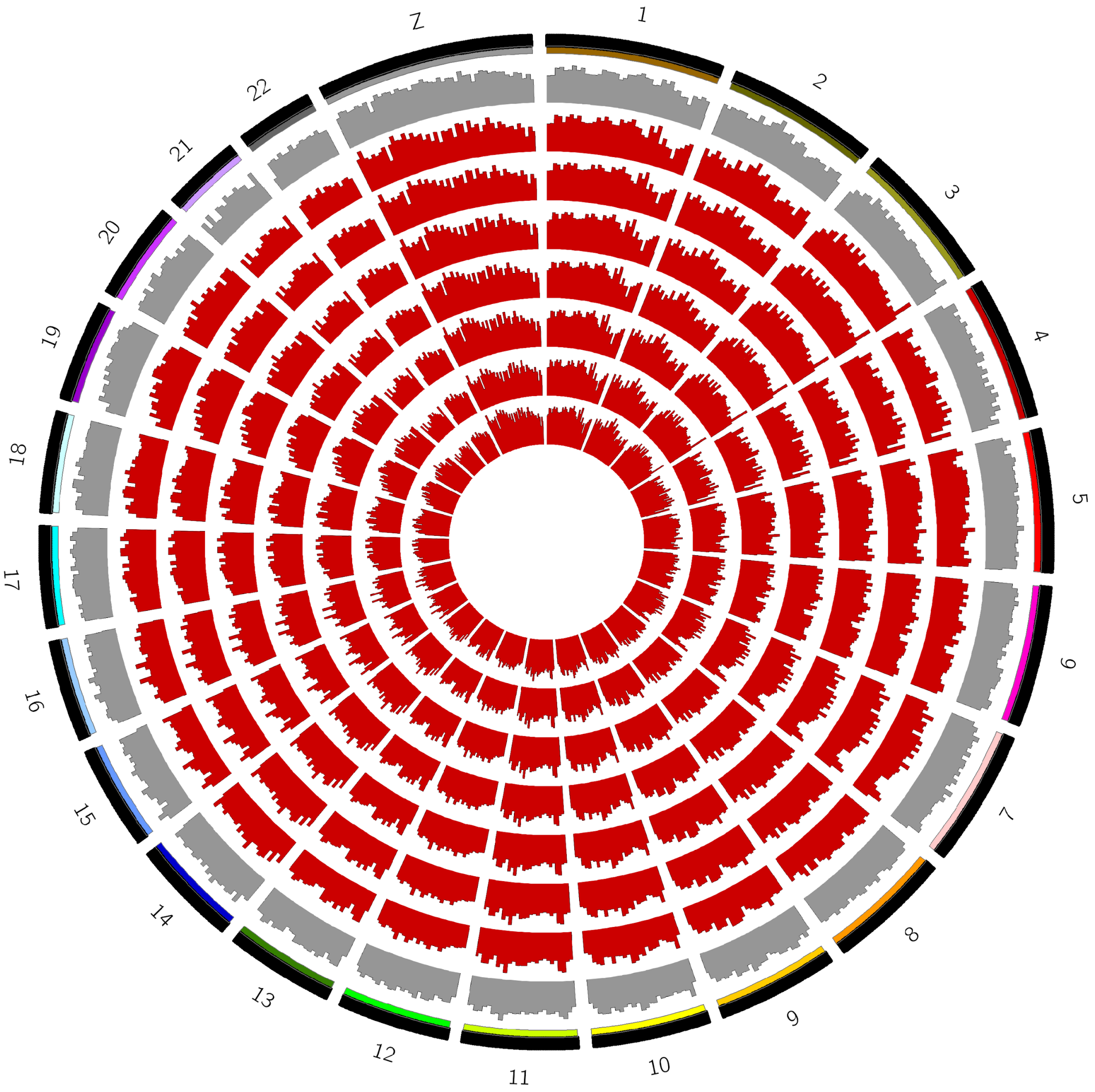


**Fig. S5: Coverage distribution of *G. lygdamus* samples across the reference.** Reference mappability is represented by the outermost grey histogram. Red histograms represent the coverage of *G. lygdamus* samples across the *G. alexis* reference in windows of 1mbp. The sample represented are, from outer to inner rings: L002, L004, L006, L008, L011, L012 and L013.

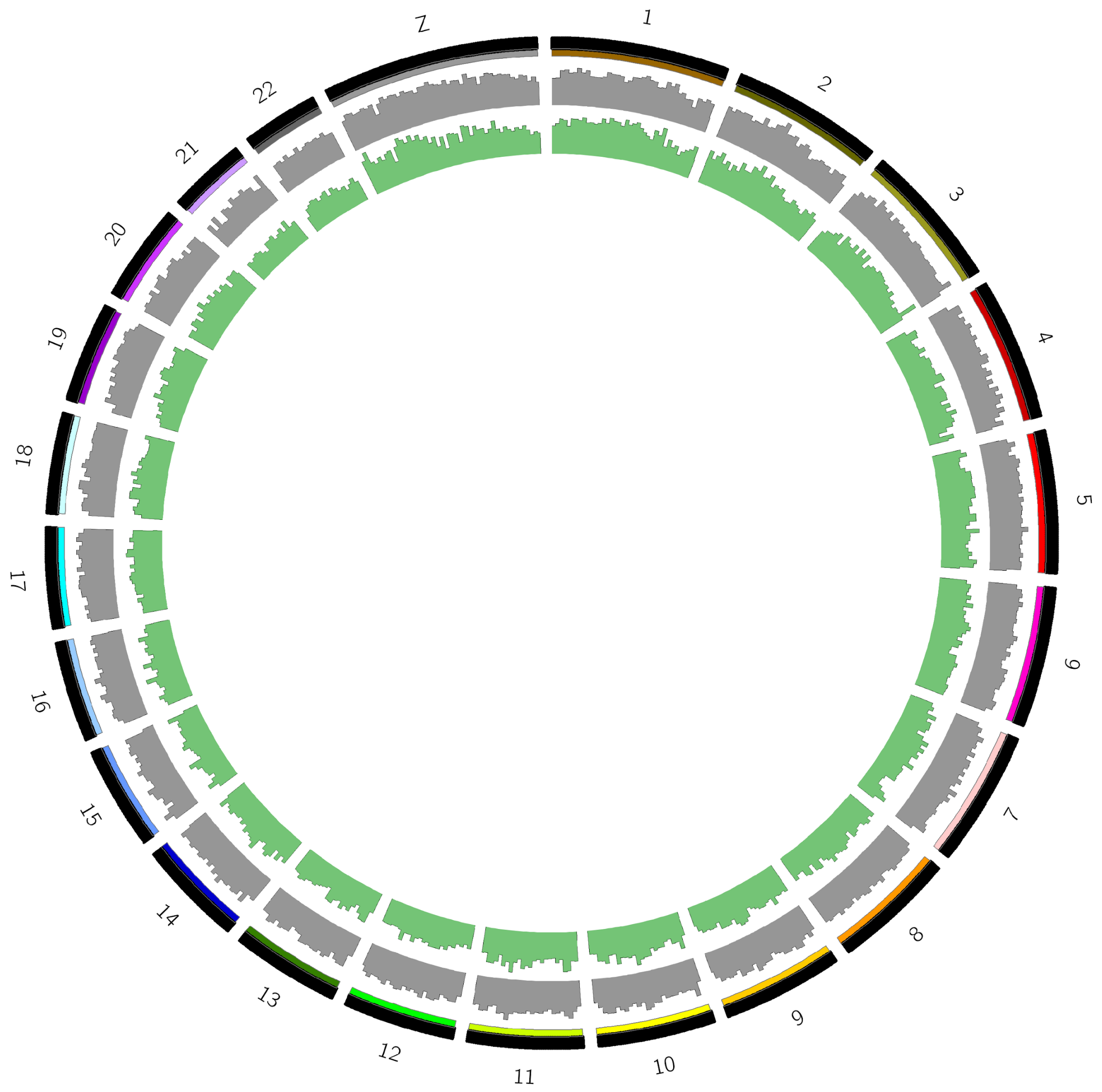


**Fig. S6: Coverage distribution of *G. lygdamus* samples across the reference.** Reference mappability is represented by the outermost grey histogram. Green histograms represent the coverage of *G. lygdamus* samples across the *G. alexis* reference in windows of 1mbp. The sample represented are, from outer to inner rings: RVcoll10-B005 (Canadian specimen).

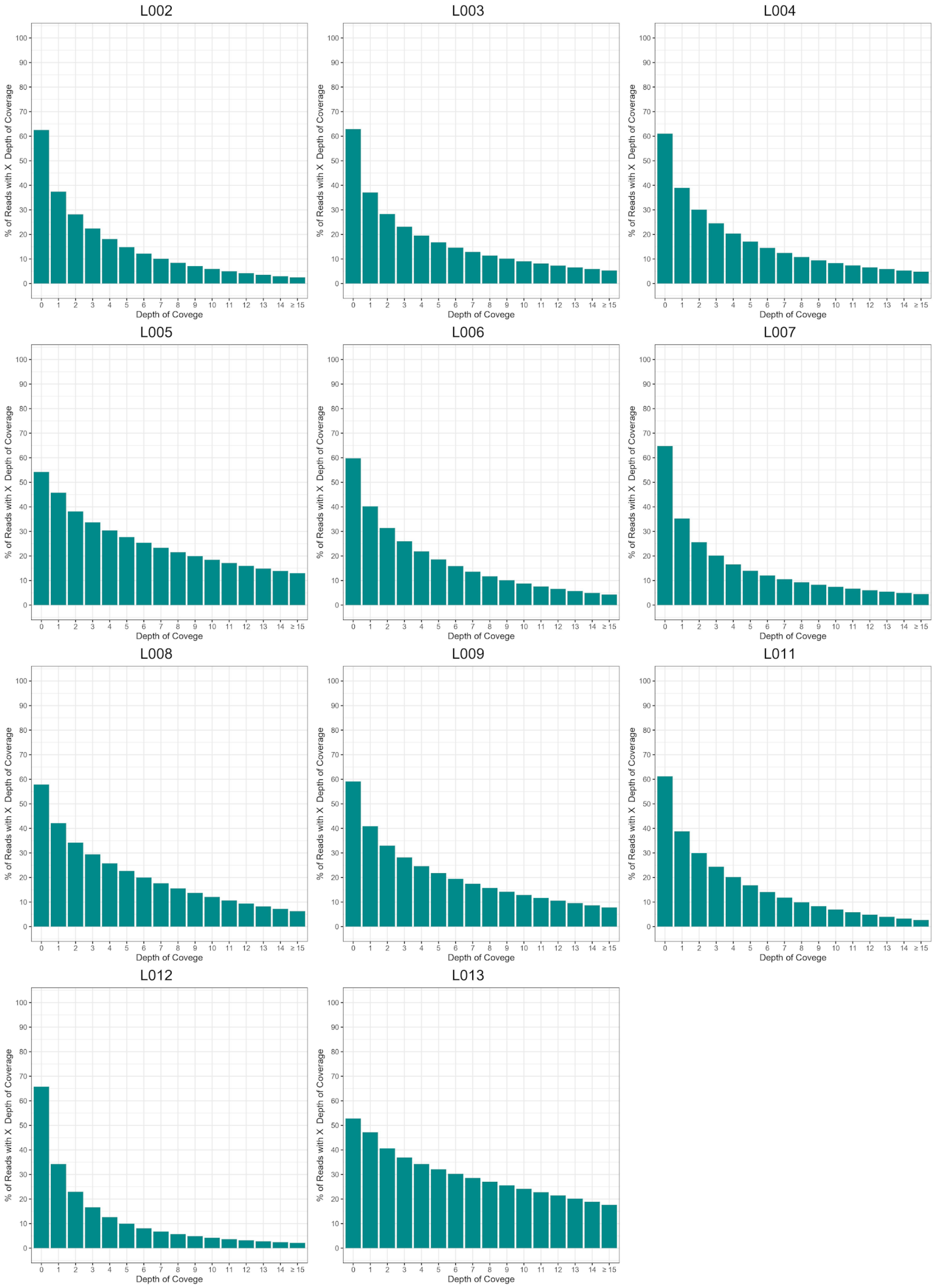


**Fig. S7: Depth of coverage distribution per position of the historical samples.**

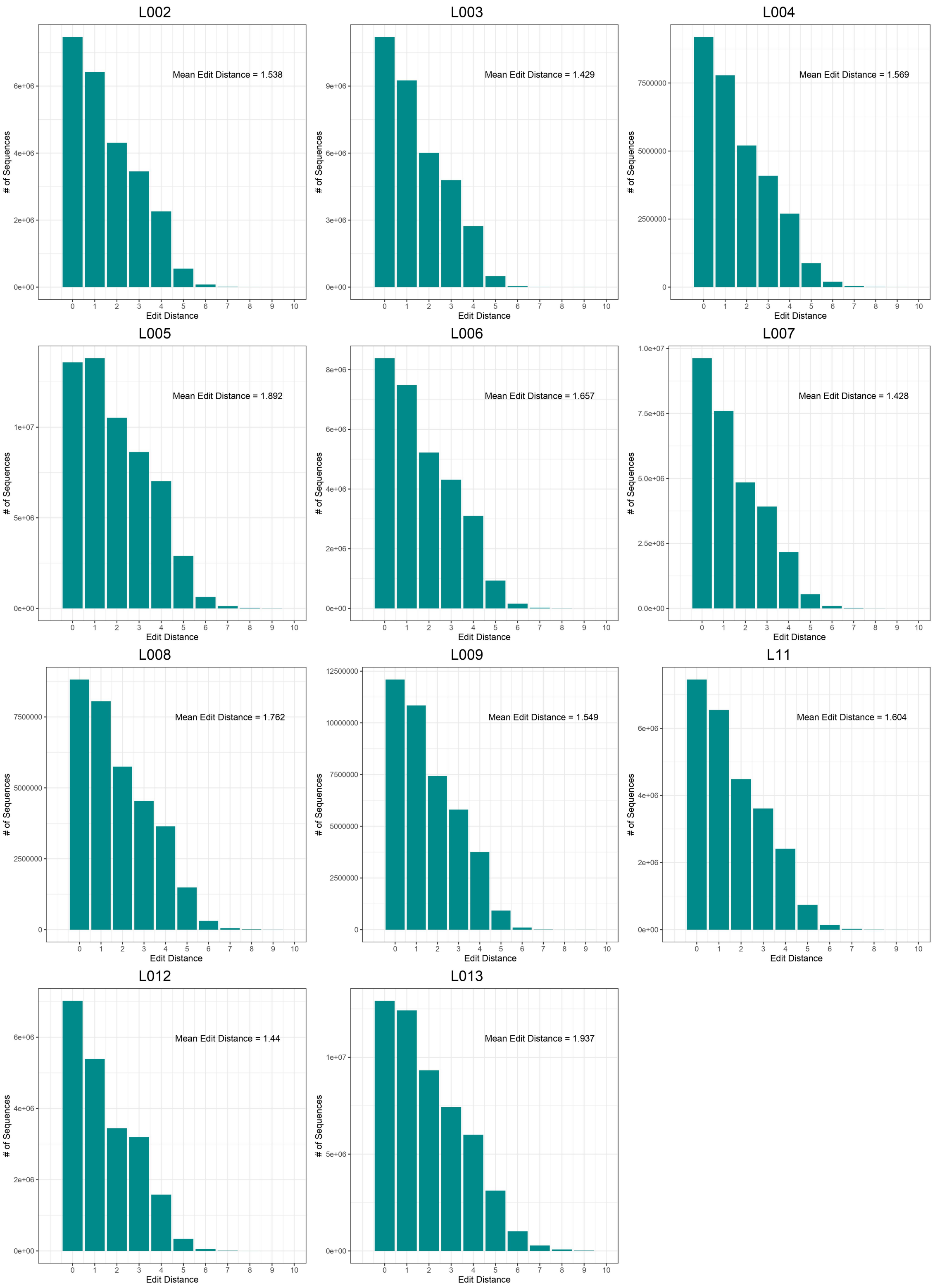


**Fig. S8: Mapped reads edit distance distribution across the 11 analysed historical samples.**

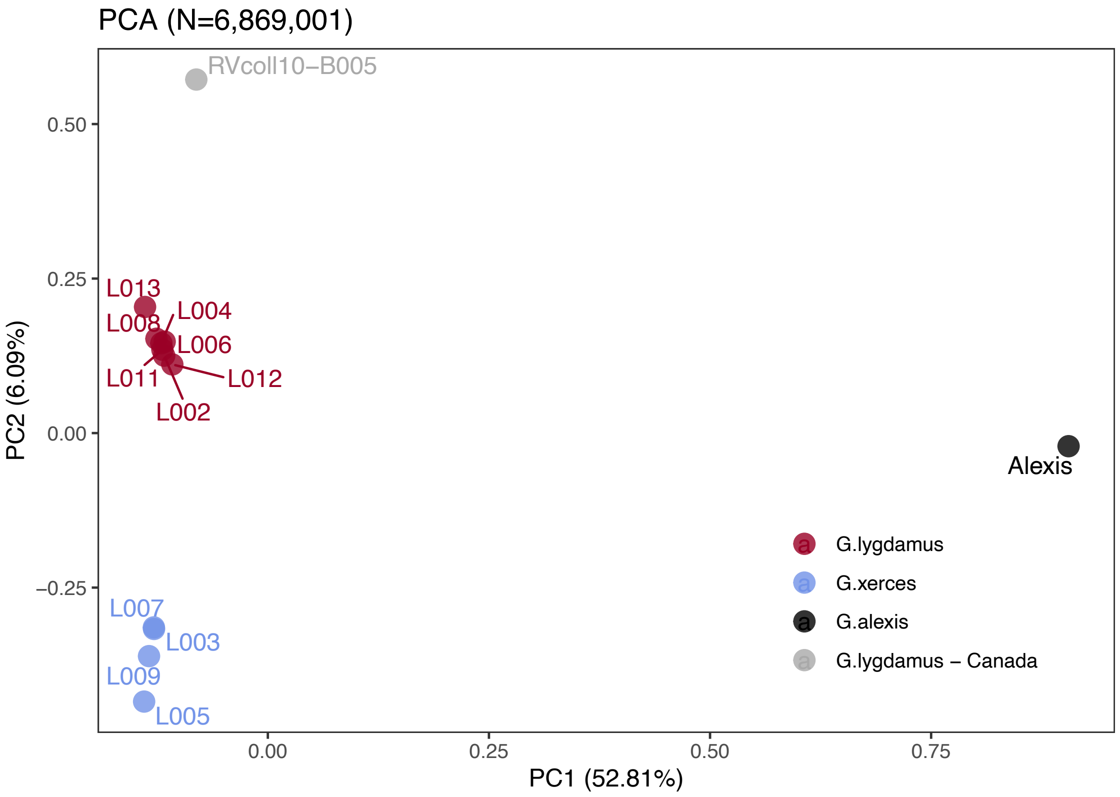


**Fig. S9: Plotting of PC1 and PC2 of the Principal Component Analysis (PCA).** The PCA was generated with nuclear DNA data (N=6,682,591 SNPs) from eleven historical butterfly specimens (4 *G. xerces* and 7 *G. lygdamus*), a modern *G. lygdamus* from Canada (RVcoll10-B005) and a modern *G. alexis* reference genome. The PCA shows a clear separation of both historical species and the reference in the first PC (explaining 52.81% of the variance), and separation of *G. xerces* and *G. lygdamus* by the second PC (explaining 6.09% of the variance), supporting they are separated lineages.

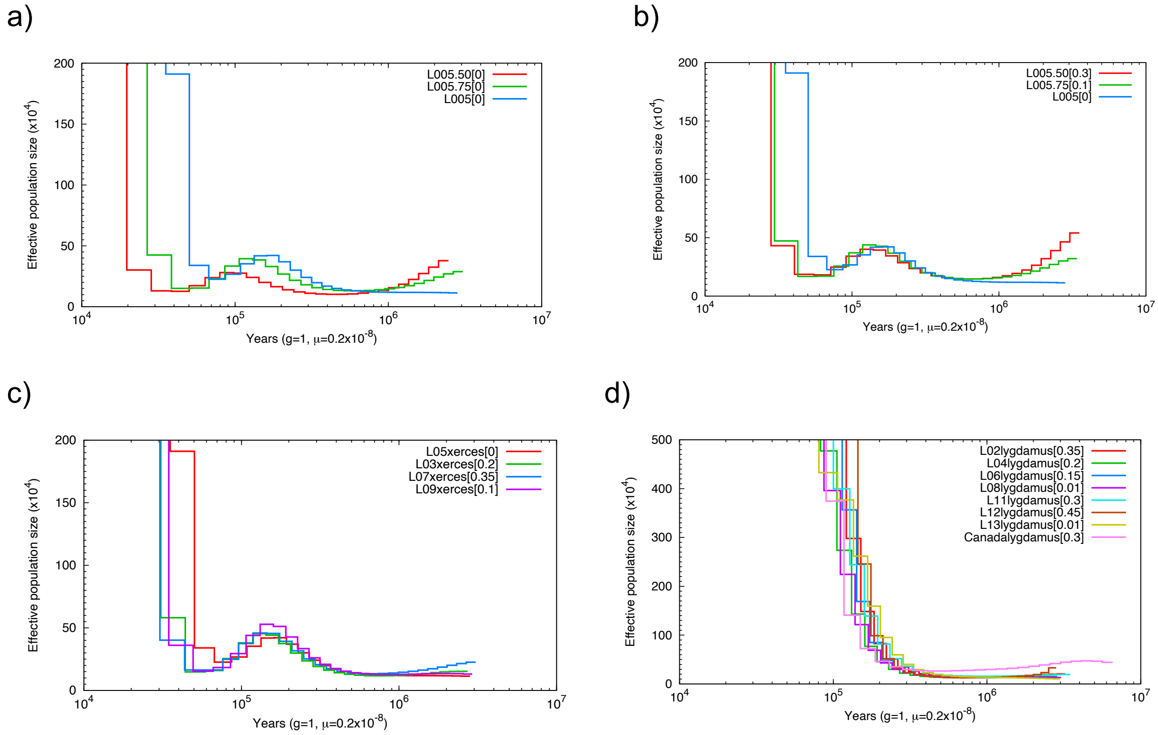


**Fig. S10: Pairwise Sequentially Makovian Coalescent (PSMC) plots of Xerces Blue and Silvery Blue.** a): PSMC of Xerces Blue L05 down sampled to half (red) and 75% of coverage (green). b) PSMC of down sampled Xerces Blue L05 corrected. Lower coverage results in underestimation of heterozygote site and thus lower historical effective population sizes. This situation can be corrected assuming a False Negative Rate (FNR) by visually adjusting the curves using the psmc_plot.py program from the PSMC package. c) PSMC of Xerces Blue L03, L05, L07 and L09 corrected assuming FNR. d) PSMC of historical Silvery Blue L02, L04, L006, L08, L11, L12 and L13 and modern Silvery Blue from Canada (RVcoll10-B005) corrected assuming FNR. Despite current differences in coverage, individuals from each species follow the same trajectory.

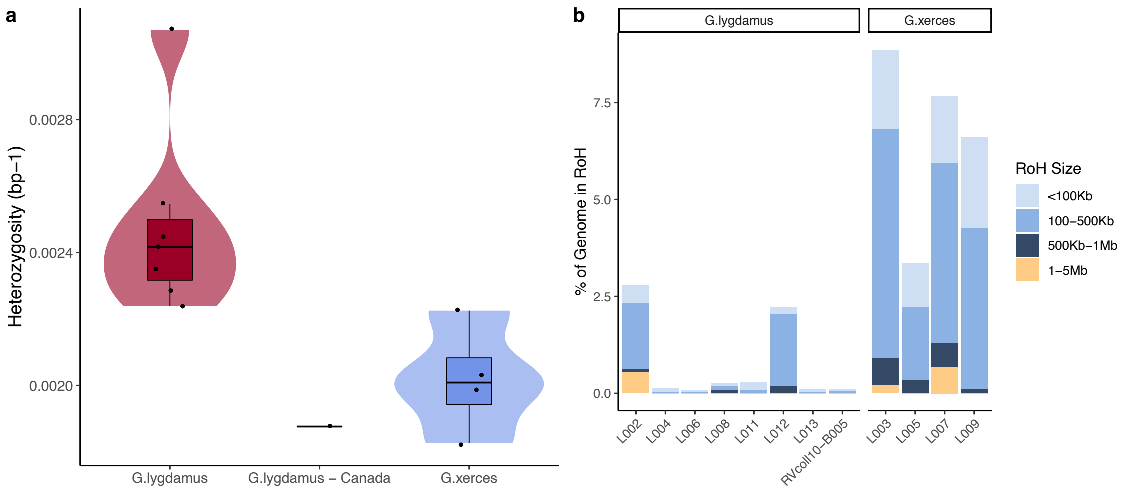


**Fig. S11: A) Heterozygosity estimates for low coverage data from both Silvery Blue (historical and modern) and Xerces Blue butterflies. B) ROHs for each individual classified by length.**

**Supplementary Tables**

**Table S1: List of historical specimens analysed in this study.**

| Genome # | Genus | Species | Subspecies | State | Locality | Date | Collection |
|---|---|---|---|---|---|---|---|
| USNMENT101413 | Glaucopsyche | xerces | | California | San Francisco | | Barnes |
| USNMENT101402 | Glaucopsyche | xerces | | California | San Francisco | April 16-23 | Barnes |
| USNMENT101441 | Glaucopsyche | xerces | | California | San Francisco | | Barnes |
| USNMENT101406 | Glaucopsyche | xerces | | California | San Francisco | | Barnes |
| USNMENT101434 | Glaucopsyche | xerces | | California | San Francisco | April 16-23 | Barnes |
| USNMENT00181297 | Glaucopsyche | lygdamus | incognitus | California | Marin County | | Barnes |
| USNMENT00181298 | Glaucopsyche | lygdamus | incognitus | California | Fairfax | 27-maig-32 | Wm D Field |
| USNMENT00181299 | Glaucopsyche | lygdamus | incognitus | California | Oakland | 14-abr-48 | Graham Heid |
| USNMENT00181300 | Glaucopsyche | lygdamus | incognitus | California | San Jose | 27-març-64 | P. Opler |
| USNMENT00181301 | Glaucopsyche | lygdamus | incognitus | California | Haywood City | 1-maig-31 | Wm D Field |
| USNMENT00181302 | Glaucopsyche | lygdamus | incognitus | California | Santa Cruz | 1-abr-32 | JW Tilden/Field |
| USNMENT00181303 | Glaucopsyche | lygdamus | incognitus | California | Santa Cruz | 8-abr-27 | GW Rawson |

**Table S2: Mapping Statistics, Mappability and Heterozygosity-RoH (Supplementary File).**

**Table S3: Mitochondrial genomes used for phylogenetic analysis.**

| Specie | Assembly ID |
|---|---|
| *Aricia agestis* | LR990279.1 |
| *Aricia artaxerxes* | OW569311.1 |
| *Celastrina argiolus* | LR994603.1 |
| *Cyaniris semiargus* | LR994570.1 |
| *Glaucopsyche alexis,* | FR990065.1 |
| *Glaucopsyche xerces* | MW677564.1 |
| *Lysandra bellargus* | HG995365.1 |
| *Lysandra coridon* | HG992145.1 |
| *Plebejus argus* | MN974526.1 |
| *Plebejus melissa* | DWQ001000057.1 |
| *Plebejus anna* | DWTA01000073.1 |
| *Polyommatus icarus* | OW569343.1 |
| *Shijimiaeoides divina* | NC_029763.1 |
| *Zizina emelina* | MN013031.1 |

**Table S4: Genes in unrecoverable regions (Supplementary File).**

**Table S5: Coordinates of the analysed coloration genes.**

| Chromsome | Start | End | Gene |
|---|---|---|---|
| FR990043.1 | 5387706 | 5403599 | Wnt1 |
| FR990043.1 | 5417902 | 5423677 | Wnt6 |
| FR990043.1 | 5519353 | 5539737 | Wnt10b |
| FR990043.1 | 5553666 | 5554753 | Wnt10a |
| FR990043.1 | 26972856 | 26974487 | WntA |
| FR990046.1 | 2343467 | 2357667 | Wnt7b |
| FR990046.1 | 6255275 | 6271623 | Wnt5b |
| FR990046.1 | 19475636 | 19486554 | Wnt9 |
| FR990050.1 | 16200978 | 16212495 | Wnt11 |
| FR990054.1 | 20633400 | 20655261 | Cortex |
| FR990059.1 | 20254460 | 20255275 | Optix |

**Table S6: Wolbachia reads assigned using Kraken2.**

| Specimen | Wolbachia Genera Reads | Wolbachia Spp. Reads |
|---|---|---|
| L002 | 190 | 5 |
| L003 | 131 | 3 |
| L004 | 213 | 5 |
| L005 | 311 | 8 |
| L006 | 242 | 9 |
| L007 | 152 | 2 |
| L008 | 414 | 21 |
| L009 | 236 | 6 |
| L011 | 184 | 9 |
| L012 | 168 | 9 |
| L013 | 523 | 24 |
